# Plasma pNfH differentiate SBMA from ALS

**DOI:** 10.1101/549576

**Authors:** Vittoria Lombardi, Bombaci Alessandro, Luca Zampedri, Ching-Hua Lu, Bilal Malik, Henrik Zetterberg, Amanda Heslegrave, Carlo Rinaldi, Linda Greensmith, Michael Hanna, Andrea Malaspina, Pietro Fratta

**Author notes:** Address for correspondence: Dr Pietro Fratta University College London Institute of Neurology, Queen Square, London, WC1N 3BG, UK. and Dr Andrea Malaspina Queen Mary University 4 Newark St, London, E1 2AT, UK. These authors contributed equally to this work (Vittoria Lombardi, Bombaci Alessandro).

## Abstract

**Background and aim:** Spinal bulbar muscular atrophy (SBMA) is a progressive adult-onset X-linked neuromuscular disease. Although traditionally considered a motor neuron disorder, recent advances have highlighted a primary myopathic component. We evaluated levels of phosphorylated neurofilament heavy chain (pNfH), a known biomarker for neurodegeneration, in SBMA.

**Materials and methods:** We collected plasma and serum from 46 SBMA, 50 ALS and 50 healthy control cases, alongside with plasma from a mouse model of SBMA (AR100) and littermate controls. We measured pNfH plasma levels using Single molecule array (Simoa), we assessed functional scales and we gathered demographic data. We analysed data using Mann-Whitney U test, Kruskal-Wallis test and Cox regression analysis.

**Results:** Plasma pNfH levels were significantly increased in ALS, but, intriguingly, there was no change in SBMA. These results were also confirmed in SBMA mice. The ROC curve highlighted that pNfH levels can effectively distinguish between ALS and SBMA (AUC 0.95).

**Conclusions:** Unexpectedly, levels of pNfH are normal in SBMA, whilst they are increased in ALS, and suggest pNfH could serve as a biomarker to differentiate the two diseases. Further, this finding is in agreement with recent evidence showing that primary muscle damage is a crucial feature in SBMA.

## Introduction

Spinal and bulbar muscular atrophy (SBMA), also known as Kennedy disease (KD), is a slowly progressive adult-onset X-linked neuromuscular disorder any effective treatments. It is characterized by limb and bulbar muscle wasting, weakness and fasciculation, associated with metabolic and endocrine alterations^1,2^. SBMA is caused by the expansion of a CAG repeat in exon 1 of the androgen receptor (*AR*) gene; more than 37 repeats are pathogenic^1^.

There is a strong need for disease activity and progression markers, which are necessary for effective clinical trials. Furthermore, although the genetic test is diagnostic, there is a lack of biomarkers to aid the initial differential diagnosis of the disease. Neurofilaments (Nfs), both light and heavy chains, are now becoming a widely accepted prognostic biomarker for amyotrophic lateral sclerosis (ALS) and other neurodegenerative diseases^3^. More than twenty years ago neurofilaments were first found to be increased in ALS CSF^4^. Reliable measurements of Nfs in peripheral biofluids have become available over the last five years, NfL were found to predict disease course in ALS and both NfL and pNfH are increased in a wide range of neurological conditions^3,5,6^. Nfs are currently being used as a biomarker in numerous clinical trials.

Recently, plasma NfL levels were found to be normal in SBMA patients^14^. This finding has supported other lines of evidence that point to a primary myopathic component in SBMA^2,7,8^.

We here investigated pNfH plasma levels in SBMA patients and in a well-established rodent model of disease, using the highly sensitive Simoa platform.

## Materials and Methods

### Study design

This is a cross-sectional study in which we enrolled SBMA and ALS patients, visited in Motor neuron clinics at University College of London Hospital and at Queen Mary Hospital between Sep 2009 and Nov 2017. We included 46 SBMA patients, 50 ALS patients and 50 healthy controls (HCs). ALS were subdivided in 2 sub-groups (ALS-Fast and ALS-Slow) on the basis of their disease progression rate to last visit (PRL): PRL ALS-Slow<0.6 and ALS-Fast >0.9. PRL was calculated as 48 (corresponding to the in-healthy state, before symptoms onset) minus the ALS Functional-Rating-Scale revised score (ALSFRSr) at the last visit, divided by time interval in months between symptoms onset and last visit date.

We collected plasma after patients’ written informed consent. Approvals were obtained from the East London and the City Research Ethics Committee (09/H0703/27).

We also assessed the test in mice, 10 well-established mouse models of SBMA (AR100) and 10 wild type littermate controls.

Samples were processed, stored and analysed as previously described^5^. All blood samples were collected into EDTA-containing tubes, centrifuged at 20°C at 3.500 rpm for 10 minutes within 1 hour and stored at −80°C.

We measured pNfH levels in plasma performing a single molecule array (Simoa)-based assay (Quanterix, Lexington, MA)^9^, using pNF-Heavy Discovery Kit 102669.

### Clinical assessment

SBMA patients had a genetical confirm of diagnosis, while ALS patients had a diagnosis of definite or probable ALS according to the revised El-Escorial criteria^10^. Disease severity was assessed using ALSFRSr^11^, SBMA Functional-Rating-Scale (SBMAFRS)^12^ and Adult-Myositis-Assessment-Tool (AMAT)^13^ scale in SBMA patients and using ALSFRS-r scale in ALS patients. Demographic and clinical data of patients are gathered in Table 1.

**Table 1 –.** Cohort demographic and genetic information, pNfH and scales values

| Patients Group | n° | Sex M/F | Age ± SD (y) | n° CAG ± SD | PRL ± SEM | pNfH ± SEM (pg/ml) | ALSFRS<sub>r</sub> ± SEM | SBMAFRS ± SEM | AMAT ± SEM |
| --- | --- | --- | --- | --- | --- | --- | --- | --- | --- |
| SBMA | 46 | 46/0 | 56,6 ± 11,1 | 42.1 ± 0.4 | N/A | 42 ± 4.9 | 38.8 ± 0.8 (n=37) | 41.1 ± 1.3 (n=36) | 31.8 ± 1.4 (n=38) |
| ALS-Slow | 25 | 14/11 | 66,9 ± 12,7 | N/A | 0.27 ± 0.04 | 629 ± 159 | 37.6 ± 1.6 | N/A | N/A |
| ALS-Fast | 25 | 10/15 | 66.4 ± 10.7 | N/A | 1.61 ± 0.12 | 1615 ± 327 | 31.7 ± 1.8 | N/A | N/A |
| HC | 50 | 17/33 | 58,5 (6,8) | N/A | N/A | 76 ± 12.7 | N/A | N/A | N/A |

Abbreviations: Healthy controls (HCS), spinal bulbar muscular atrophy (SBMA); fast progressing amyotrophic lateral sclerosis (ALS-Fast); slow progressing amyotrophic lateral sclerosis (ALS-Slow); standard deviation (SD), standard error of the mean (SEM); years (y); female (F); male (M); number (n°); number of CAG repeats in androgen receptor gene (n° CAG); adult myopathy assessment tool (AMAT) ; spinal bulbar muscular atrophy function rating scale (SBMAFRS); amyotrophic lateral sclerosis function rating revised scale (ALSFRS<sub>r</sub>); phosphorylated neurofilament heavy chain (pNfH); progression rate to last visit (PRL).

### Statistical analysis

Mann-Whitney U test and Kruskal-Wallis tests were performed to analyse plasma pNfH levels between groups. Dunn’s multiple comparisons test was performed following Kruskal-Wallis test in case of significant differences. Receiver operating characteristic (ROC) curve, and the corresponding sensitivity, specificity, positive and negative predictive values, accompanied by their 95% CIs, was performed in order to identify the best cut-off level of pNfH to separate ALS and SBMA patients. Correlation between parameters was calculated by Spearman rank correlation r. The level of significance for all statistical tests was set at 0.05. The program Prism V.8 (GraphPad Software, La Jolla, CA) was used to perform statistical calculations.

## Results

This study included 46 patients with SBMA, 50 ALS patients and 50 HCs. Participant demographic and clinical characteristics are summarized in Table 1. We measured pNfH on plasma samples using a Simoa assay and found levels to be unchanged in SBMA vs HCs (**Figure 1A**). Of note, mean pNfH levels in SMBA were lower than HCs, although this difference was not statistically significant using Dunn’s multiple comparison test. These results were also confirmed in SBMA mice, where pNfH levels were lower than WT mice (p= 0,009) (**Figure 1B**). Conversely, in both fast-and slow-progressing ALS groups there was a statistically significant increase of plasma pNfH levels compared to HCs (ALS-Slow p<0.001; ALS-Fast p<0.0001), conforming with previous reports^3,5^. The two groups significantly differed when performing a Mann-Whitney test (p=0.012), although significance was not retained after Dunn’s multiple comparison correction. Therefore, in order to further investigate whether pNfH levels are associated to disease severity in ALS, we tested the correlation between pNfH levels and disease progression rate and observed a moderately significant association (r_s_=0.36, p=0.01).

**Figure 1.**
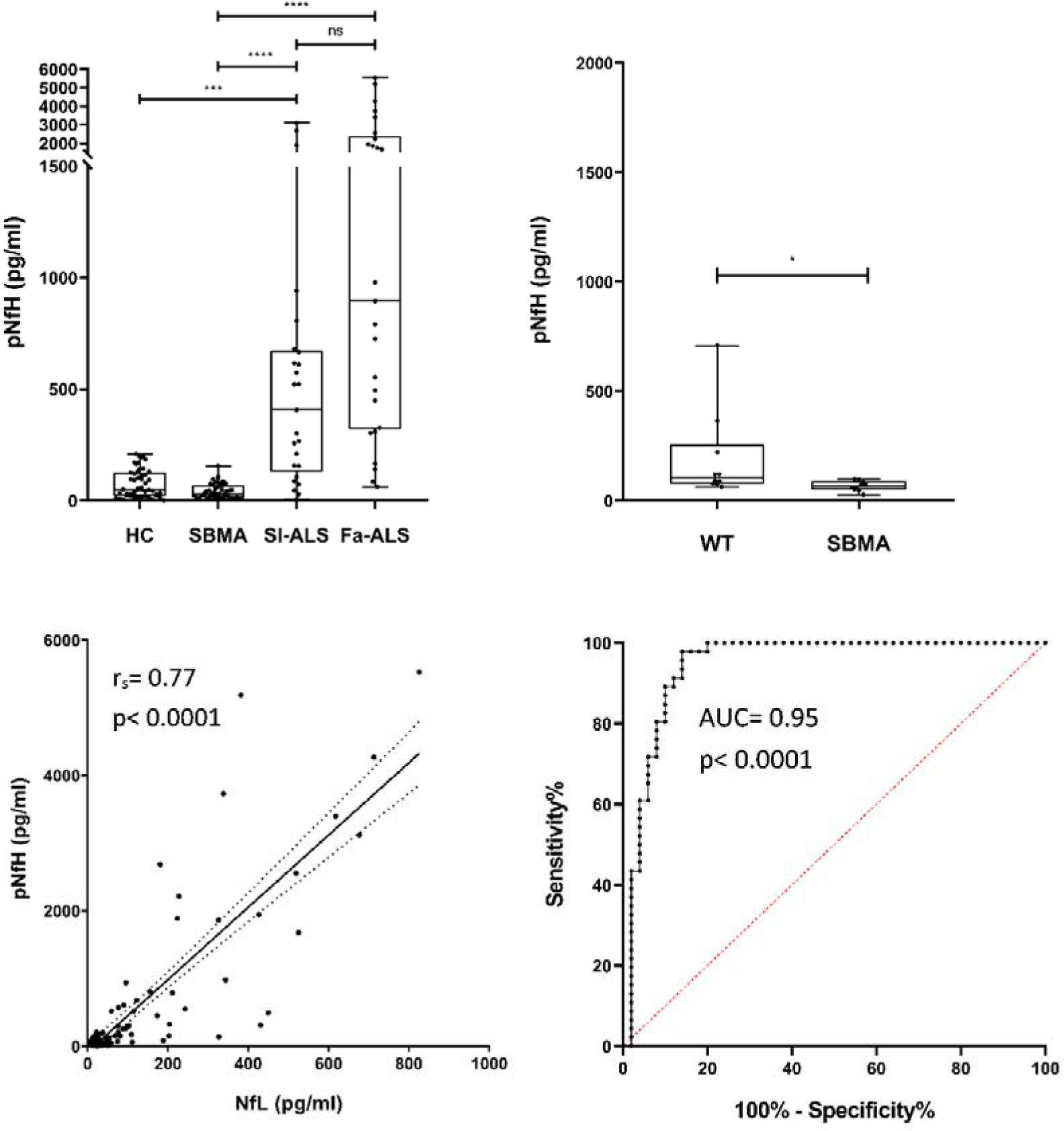
(**A**) pNfH concentration in HCs, SBMA, ALS-Slow and ALS-Fast. Using Kruskal-Wallis test and Dunn’s multiple comparison test we observed reduced levels of both pNfH compared to ALS groups (all p-value < 0.0001). (**B**) pNfH levels from mice AR100 (SBMA) and littermate controls (WT) show a light statistically significant difference (Mann Whitney test). (**C**) Correlation between pNfH and NfL (r_s_= 0.77, p< 0.0001). (**D**) ROC curve, created comparing pNfH and NfL levels, shows a high AUC (0.95; 95% CI 0.90-1.00). The value associated to the highest Youden-Index is 105 pg/ml (98% sensitivity, 86% specificity, PPV 0.88, NPV 0.98). Fa-ALS= Fast-progressing amyotrophic lateral sclerosis; Sl-ALS= slow-progressing amyotrophic lateral sclerosis; WT= wild type mice; HCs= Healthy controls; SBMA= spinal bulbar muscular atrophy; pNfH= phosphorylated-Heavy-Neurofilaments; NfL= Light-Chain-Neurofilaments; AUC= area under curve; PPV= positive predictive value: NPV= negative predictive value; p= p-value. ns= not statistically significant result; *p< 0.05; **p< 0.01; ***p< 0.001; ****p< 0.0001.

We then tested whether pNfH levels correlate with disease functional impairment. No correlation was found between pNfH and ALSFRSr, SBMAFRS and AMAT scales in SMBA patients (r_s_=-0.029, p=0.86; r_s_=0.025, p=0.89; r_s_=0.024, p=0.89, respectively) or between pNfH and ALSFRSr in ALS patients (r_s_=-0.193, p=0.18). pNfH did not correlate with age in all groups.

Importantly, pNfH plasma levels were significantly different between SBMA and both ALS groups (p<0.0001). In order to identify the pNfH threshold to most effectively distinguish SBMA from ALS, we created a ROC curve (AUC=0.95, p<0.0001; **Figure 1D**) and defined pNfH of 105 pg/ml (highest Youden-Index) as the best cut-off between ALS and SBMA (98% sensitivity, 86% specificity). Given the similarity of results previously reported with NfL, we compared pNfH with previously measured NfL levels, in the same samples^14^. Cox regression model analysis, showed a significant correlation between pNfH and NfL levels (r_s_=0.77, p<0.0001; **Figure 1C**).

## Discussion

Contrary to what would be expected in a motor cell disorder^3,14,15^, this study shows low levels of pNfH in SBMA patients rather than a disease-related increase, as demonstrated in ALS and in primary lateral sclerosis. The result was also confirmed in a well-established SBMA mouse model.

We highlighted an increase of pNfH levels in ALS, and, beyond to what previously reported^15^, we have also found a correlation between pNfH levels and disease progression rate in ALS (calculated as PRL). As pNfH levels are increased in the initial phases of ALS^15^, this finding could be complementary to genetics to provide an early differential diagnosis for SBMA and other lower motor neuron diseases. It could also bear importance in predicting disease progression and in patient stratification for clinical trials.

We have also seen no relationships between pNfH levels and the stage of disease (quantified using functional and motor scales) both in SBMA and in ALS patients. This could suggest an absence of variation of pNfH levels during the course of these diseases, confirming previous data obtained in ALS with in-house ELISA test^15^.

Moreover, pNfH and NfL measurements in the same patients showed a good correlation; this correlation is tighter in ALS group than SBMA and HCs groups, due to the highest variability of measurement when detecting low levels of neurofilaments. Further work will be needed to assess whether combining these measurements could ameliorate the predictive value of the test.

Using ROC curve, we were also able to identify a statistically strong cut-off level (AUC=0.95) of pNfH that could distinguish SBMA from ALS. This suggests that pNfH could be used as a biomarker for the differential diagnosis between SBMA and ALS.

Limits of this study are represented by the cross-sectional design that does not allow us to infer about the variation of pNfH during the disease course. With regard to the analytical aspect, in this study we employed one of the most sensitive platforms for neurofilament analysis; this approach was the same used for the NfL study^14^. Unlike the measurement of NfL, previous work on the detection of plasma neurofilament isoforms has highlighted the inherent difficulties encountered in pNfH measuring in biological fluids, including the lack of linearity in dilution experiments and the presence of pNfH immunocomplexes or aggregates in circulation; that could cause epitope masking and make measurements for clinical stratification difficult^15^.

Finally, normal levels of pNfH in SBMA are in contrast not only with ALS, but also with previous findings on peripheral neuropathies^3,5^, that share a slow disease course, although have a mild to moderate pNfH increase. Unfortunately, the latter data resulted from small cohorts of patients, included in largest studies, or from case reports and pNfH was measured using in-house ELISA test; so there could have been an underestimation of results.

This data supports the recent hypothesis of a primary muscle origin of damage in SBMA patients^2,4,13,14^ and is consistent with results of NfL in SBMA^14^.

Under this light, it could be useful to measure, in a prospective study, pNfH levels in primary muscle atrophy and other pure lower motor neuron diseases, using Simoa platform.

## Conclusions

In conclusion, we have demonstrated that plasma pNfH concentrations are not increased in SBMA patients and in a mouse model of disease, as opposed to ALS, and that pNfH could be used as a biomarker in differential diagnosis between SBMA and ALS.

## Acknowledgements

We thank the patients involved in the study and their families for the participation and support to KD research. All samples were obtained from the ALS Biomarker study. PF is supported by an MRC/MNDA LEW Clinician Scientist Fellowship and the NIHR UCLH BRC. This study was supported by the NIHR UCLH BRC, the UCL Kennedy’s Disease Fund and KDUK. OJZ is funded by the UK National Institute of Health Research on an Academic Clinical Fellowship. CR is funded by a Welcome Trust Clinical Research Career Development Fellowship and the Muscular Dystrophy Association (MDA). HZ is funded by the UK Dementia Research Institute at UCL, the European Research Council, the Swedish Research Council and the Knut and Alice Wallenberg Foundation. The Simoa instrument was bought using a Welcome Trust multi-user equipment grant (PI: HZ).

